# Teaser: Individualized benchmarking and optimization of read mapping results for NGS data

**DOI:** 10.1101/025858

**Authors:** Moritz Smolka, Philipp Rescheneder, Michael C. Schatz, Arndt von Haeseler, Fritz J. Sedlazeck

**Affiliations:** Center for Integrative Bioinformatics Vienna, Max F. Perutz Laboratories, University of Vienna, Medical University of Vienna, A-1030 Vienna, Austria; Bioinformatics and Computational Biology, Faculty of Computer Science, University of Vienna, Vienna, Austria; Simons Center for Quantitative Biology, Cold Spring Harbor Laboratory, Cold Spring Harbor, NY

**Keywords:** Mapping, benchmarking, individualized data, Next Generation Sequencing, High throughput sequencing

## Abstract

Mapping reads to a genome remains challenging, especially for non-model organisms with poorer quality assemblies, or for organisms with higher rates of mutations. While most research has focused on speeding up the mapping process, little attention has been paid to optimize the choice of mapper and parameters for a user’s dataset. Here we present Teaser, which assists in these choices through rapid automated benchmarking of different mappers and parameter settings for individualized data. Within minutes, Teaser completes a quantitative evaluation of an ensemble of mapping algorithms and parameters. Using Teaser, we demonstrate how Bowtie2 can be optimized for different data.

## Rationale

Recent and ongoing advances in sequencing technologies and applications [1, 2] lead to a rapid growth of methods that align next generation sequencing (NGS) reads to a reference genome (read mapping). By mid 2015, nearly 100 different mappers are available, although not all are equally suited for a given application or dataset [3]. The large number of potential options, and the even larger number of potential parameter configurations makes it challenging to choose the most appropriate mapper for a given dataset. Consequently most users generally rely on the default, unoptimized, parameters for one of a few popular algorithms, even when this choice performs very poorly compared to an optimized approach. This may introduce substantial biases in the subsequent analysis, including reduced coverage, reduced rates of mutations or heterozygosity, false determination of allele-specific expression, or other artifacts [4, 5].

Previous research [3, 6-8] has focused on benchmarking mappers for particular scenarios (e.g. SNP calling, or data from a specific sequencing instrument) or for selected organisms. Although these surveys represent a valuable resource for certain tasks, they are most often performed using only the default parameters and versions of the software that can be outdated by the time they are published. More significantly, these evaluations may not capture the data types or genomes used in the study at hand, which may have substantially different characteristics. To choose the most suitable parameter settings for a given mapper requires in depth knowledge of the data as well as the mapper. This is extremely complex to do, and in many cases, even the author(s) of the software may not fully appreciate how to best optimize their own software for a given dataset.

Recent efforts to provide guidance for choosing the mapper and its parameter setting like GCAT focus on human data [8]. GCAT is an online resource that hosts simulated reads for a version of the human genome that users can download and analyze, using their own analysis pipelines. Afterwards the results can be uploaded and are compared to the gold standard. On a voluntary basis the parameter settings of the analysis are made publicly available for the benefit of the community. However, not all researcher work with the human genome, and instead many researchers face the challenge of markedly different genome and read characteristics including the SNP rate, error rate, read lengths, quality of the reference genome, and reference sequence complexity such as GC content and repetitive regions, all of which influence the ability of mappers to align reads [3]. For example, whereas some mappers efficiently map reads to a human reference genome, they might be less adequate when applied to a draft *de-novo* assembled genome with incomplete and/or fragmented contigs. In any case, the choice of the mapper parameters depends on the characteristics of the data.

Here we introduce Teaser, a method that assists users to determine the most suitable mapper and parameters given the core characteristics of their individual experiment. Teaser simulates read data, executes a number of popular mapping tools under an ensemble of parameter settings, and then evaluates and illustrates the results. Teaser can also be used to optimize the mapping of genuine NGS reads, especially to account for difficult to simulate characteristics such as run-specific error modes or sequencing biases. Teaser‘s short runtime enables users to evaluate a multitude of different scenarios. Furthermore, Teaser is highly flexible, easily allowing to: (i) extend the catalog of mappers, (ii) customize mapper parameters, (iii) provide your own simulation or select from a list of preconfigured simulation methods, and (iv) fine-tune the evaluation of mappers. Teaser provides several summary statistics from the experiments such as the fraction of correctly/wrongly mapped reads, correctly mapped reads per second, precision and recall, F-measure (i.e. the harmonic mean of precision and recall)[7], maximum memory usage and runtime. In the end, Teaser generates a HTML based report including interactive figures that can be viewed using common web browsers. Teaser is available open-source as a web application (teaser.cibiv.univie.ac.at), a virtual machine image, and as a standalone version (github.com/Cibiv/Teaser).

## Teaser description and data characteristics

Teaser comprises three main steps. First, it simulates reads based on user-defined specifications (e.g. sequencing technology, sequencing error model, read length, SNP rate) and a reference genome. Second, Teaser automatically executes the mappers, monitors and evaluates their performance. Third, it summarizes the evaluation results and generates a HTML based report. In the following we will describe each step.

### Simulation of reads from subsampled reference genomes

The first stage of the simulation encompasses randomly subsampling regions from the reference genome. By default, the length of each region is tenfold of the user specified average read length or insert size in the case of paired-end sequencing. We introduce the following default sampling-rates based on genome size. Sampling stops if the total length of sampled regions exceeds 50% of the reference genome length for small (<100mb) or 25% of medium sized genomes (<500mb). For larger genomes (e.g. human) Teaser samples regions until reaching 1% of its genome length. Overall Teaser samples at least 15mb to guarantee a robust measurement. The sampled regions are then concatenated into an artificial chromosome. To avoid reads branching from one region to another, regions are separated using two times the specified average read length or insert size of N’s as padding. Subsequently either DWGSIM [9] or Mason [10] generate simulated reads from the artificial chromosome using the user specified characteristics for read lengths, error models, and genome characteristics (rate of heterozygosity, proportion of indels, etc). Moreover, Teaser optionally accepts fastq and SAM files [11], containing user-simulated reads and their presumably correct alignment positions and strand (used for evaluation). Finally, Teaser can also accept just a fastq file of genuine or simulated reads for evaluation, although only a subset of metrics will be available since the true mapping positions are not known.

### Mapping of simulated reads

After read generation, Teaser executes the user-defined mappers with the corresponding parameter values to align the reads to the complete reference genome. By default, Teaser includes: BWA (version 0.7.12-r1039) [11], BWA-MEM (version 0.7.12-r1039) [12], BWA-SW (version 0.7.12-r1039) [13], Bowtie2 (version 2.2.5) [14] and NextGenMap (version 0.4.13) [15]. Teaser monitors the runtime and maximum memory consumption for each mapper. The reported run-times are the times needed for mapping the reads only and do not include the time for preprocessing steps (i.e. indexing the reference genome), reading in the reference sequence and initializing the mapper. Thus, these runtimes provide a useful estimation for larger read sets.

### Evaluation of simulated and real data

Teaser first checks if a mapped read exceeds the user defined mapping quality threshold (default equals 0). A mapped read with mapping quality below that threshold is counted as not mapped. Reads passing the mapping quality check are considered correctly mapped if the following conditions are true: (i) The reported starting position of the aligned read is within X bp (by default 50 percent of the defined read length) from the original starting position of the simulated read. (ii) The read is mapped to the same strand as it was simulated from. If any of these conditions is violated the read is considered wrongly mapped. In case of multi-mapping reads, only the primary alignment (the one the mapper considers to be best [11]) is taken into account. Other methods have been suggested to evaluate read alignments than comparing the distance between the correct and the reported mapping position of a read [5]. However, we argue that evaluating the distance is the most efficient and most important evaluation criteria.

Teaser further provides a way to assess the mapping rate of the mappers and parameter settings based on real data. To grant a short runtime, Teaser first subsamples the same number of reads as it would use for the simulation (see above). After the mapping, however, Teaser computes the overall percentage of mapped reads rather than the percentage of correctly mapped reads since that information is not available. We recognize that mapping additional reads does not necessarily indicate higher quality alignments, but nevertheless this information is valuable to assess the robustness of parameter choices. Other metrics, especially runtime and memory requires are recorded as before, allowing for in-depth performance optimization.

### Mapping Summary and Report

Teaser provides six statistics for further evaluation. First, Teaser outputs the number of correctly, wrongly, and not mapped reads. Next, Teaser reports the precision (fraction of correctly mapped reads compared to all mapped reads) and the recall rate (fraction of correctly mapped reads if compared to correctly mapped reads and not-mapped reads) for each mapper. Teaser computes the F-Measure, the harmonic mean of precision and recall as suggested by Caboche et al.[7]. Thus, the F score is a measure of a mapper’s accuracy, ranging from 0 (worst) to 1 (best). Finally, Teaser reports the correctly mapped reads per second, the overall runtime, and the peak memory requirement for each mapper.

All results are displayed as part of a HTML based report providing easy to read tables and interactive figures. The report allows a direct comparison of the mapping results for different summary statistics and different mapping quality thresholds. Supplementary Note 1 describes the options in more detail. We also invite the reader to visit teaser.cibiv.univie.ac.at for further details and the presentation of results.

### Parameter Optimization

In addition to benchmarking different mappers, Teaser can evaluate different parameter sets for each mapper, defined as either specific parameter values or ranges of values to be evaluated for each mapper. The second option can be used to automatically explore the impact of key parameters, such as the k-mer length, on the mapping results. If parameter ranges are defined for more than one parameter, Teaser systematically tests every combination and reports the results for each combination separately.

Finally, to identify the optimal parameter set for the user specific genomic data, Teaser provides an additional plot that shows the correctly mapped reads in percentages and the number of reads processed per second for all evaluated mapper and parameter combinations.

**Table 1.** Summary of the parameters to generate simulated reads.

| Dataset | Reference | Platform | Read Length (bp) | Library Type | Number of simulated reads | Mutation rate (%) | Seq. error rate (%) | Subsampled (%) | Source | Teaser Runtime (mins) |
| --- | --- | --- | --- | --- | --- | --- | --- | --- | --- | --- |
| H1 | GRCh37 | Illumina | 100 | paired | 313,716 | 0.1 | 0.6 | 1 | DWGSIM | 15.41 |
| H2 | GRCh37 | Illumina | 100 | single | 313,716 | 0.1 | 0.6 | 1 | DWGSIM | 11.08 |
| D1 | BDGP6 | Illumina | 150 | single | 239,544 | 0.2 | 0.6 | 25 | Mason | 5.06 |
| M1 | GRCm38 | Illumina | 150 | single | 322,561 | 0.2 | 0.6 | 1 | Mason | 19.78 |
| D2 | BDGP6 | Illumina | 150 | single | 239,544 | 5.0 | 0.6 | 25 | Mason | 5.45 |
| M2 | GRCm38 | Illumina | 150 | single | 322,561 | 5.0 | 0.6 | 1 | Mason | 19.25 |
| D3 | BDGP6 | Ion Torrent | 200 | single | 179,658 | 0.1 | 4.0 | 25 | DWGSIM | 4.18 |
| M3 | GRCm38 | Ion Torrent | 200 | single | 241,921 | 0.1 | 4.0 | 1 | DWGSIM | 20.4 |
| D4 | BDGP6 | Illumina | 22 | single | 1,633,257 | 0.2 | 0.6 | 25 | Mason | 9.76 |
| M4 | GRCm38 | Illumina | 22 | single | 2,199,284 | 0.2 | 0.6 | 1 | Mason | 58.38 |
| C1 | C. perfretum | Illumina | 100 | paired | 112,500 | 0.9 | 0.6 | 1 | Mason | 30.06 |
| C2 |  |  |  |  | 37,500 |  | 1.0 |  |  |  |

### Performance Evaluation

To demonstrate the usefulness of Teaser we benchmarked five mappers for ten simulated data sets (Table 1). Similar to the data studied by GCAT [8], we simulated two human Illumina-like read-sets assuming a genomic SNP frequency of 0.1% and a 0.03% (H1) or 0.07% (H2) probability for the occurrence of insertion and deletions. Read length was set to 100 bp, assuming a sequencing error of 2% (default of DWGSIM [9]). In addition, we simulated from the *Mus musculus* (denoted by M) and *Drosophila melanogaster* (denoted by D) genomes 150bp Illumina-like reads (D1, M1) assuming a SNP frequency of 0.07% and indel frequency of 0.03%, and to mimic a more diverse organism assuming a SNP frequency of 3.5% and indel frequency of 1.5% (D2, M2). In both cases we assumed a sequencing error of 0.6% (default of Mason [10]). Furthermore, we simulated Ion Torrent like data (D3, M3) and data with 22bp long Illumina-like reads as encountered in miRNA sequencing (D4, M4). Further details of the simulations are displayed in Table 1.

To investigate the influence of the subsampling process on the performance of the mappers we further generated five data sets with different down-sampling rates for each organism: human, *Mus musculus* and *Drosophila melanogaster*. Mason was used to simulated Illumina-like reads for the entire genome, and Teaser was applied to the full data set. Supplementary Table 2 lists the properties of each data set.

Finally, to assess the performance of the mappers on a de-novo assembly we simulated two 100bp long paired end Illumina-like data sets (C1, C2) from a *de-novo* assembly of the 1Gbp *Cottus rhenanus* genome (FJS, J.Cheng, J. <u>Altmüller, AvH, AW. Nolte</u>, under consideration). C1 and C2 were simulated with 0.9% SNP rate, to mimic a closely related population (*Cottus perifretum*) to further assess cross-species mapping performance. Based on the observation that a portion of the reads from the real data set had a higher sequencing error rate, we simulated C1 and C2 with a 0.6% (default of Mason) and 1% sequencing error, respectively. The combined data sets C1 and C2 were provided to Teaser using the import function described above.

Unless otherwise mentioned mappers were executed with their default parameters on a desktop computer with an Intel(R) Core(TM) i5-2500K (3.30GHz) quad-core CPU and 16GB of RAM. Only the data sets used to verify the downsampling process were computed on an Intel(R) Xeon(R) CPU X5650 (2.67GHz) with 32GB of RAM.

## Results

### The influence of subsampling for benchmarking mappers

Benchmarking the five mappers with Teaser based on a full human genome took more than ten hours. To reduce computing time, Teaser randomly samples non-overlapping subsequences from the genome, and from each subsequence reads were simulated as described. The simulated reads were then mapped to the entire reference genome and evaluated. Figure 1a displays the fractions of correctly mapped reads for Human, *Mus musculus* and *Drosophila melanogaster* datasets with subsampling rates ranging from 100% to 1% of the genome.

**Figure 1:**
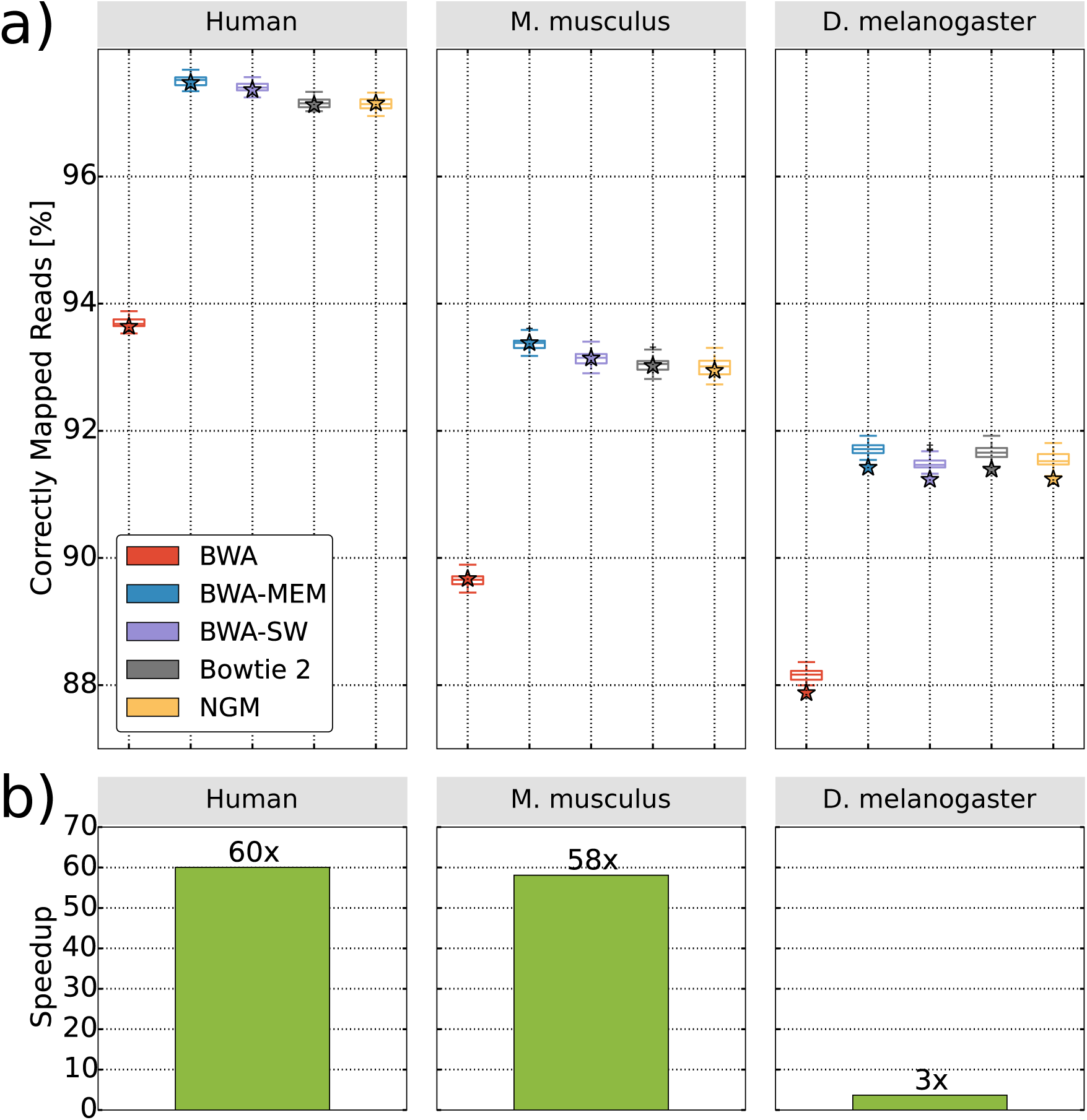
Effect of subsampling on the percentage of correctly mapped reads (a) and the runtime (b) for five mappers. For the genomes of Human, *Mus musculus* and *Drosophila melanogaster* we each generated sets of reads, once for the whole genome (shown as star symbol) and 25 times using the default subsampling rates of Teaser. The plots in (a) show, that subsampling genomic regions has an insignificant effect on the mapping rates. However, subsampling saves substantial computing time, which is most impressive for the human genome, where we observed a 60-fold reduction.

Figure 1 a shows that the proportion of correctly mapped reads is not significantly affected by the subsampling rate. The percentage of correctly mapped reads varied by less than 0.5% for the default sampling rates for the five mappers and the three reference genomes. Thus, investigating the performance of mappers on randomly sampled regions of the genome suffices. This has the great advantage that it saves computing time as shown in Figure 1b. The saving can be quite substantial, for the human genome we observe a runtime reduction from over ten hours to 13 minutes with negligible differences in observed mapping characteristics.

### Benchmarking different mappers

Teaser benchmarked the two human data sets in 26 minutes (Figure 2a, H1+H2). For both data sets, BWA-MEM outperformed the other mappers in terms of correctly mapped reads. This result is consistent with the results reported by GCAT. Furthermore, the ranking of the four mappers (NextGenMap was not evaluated) was also consistent. This shows that Teaser produced for the human data the same results as GCAT, although Teaser has much greater flexibility.

**Figure 2:**
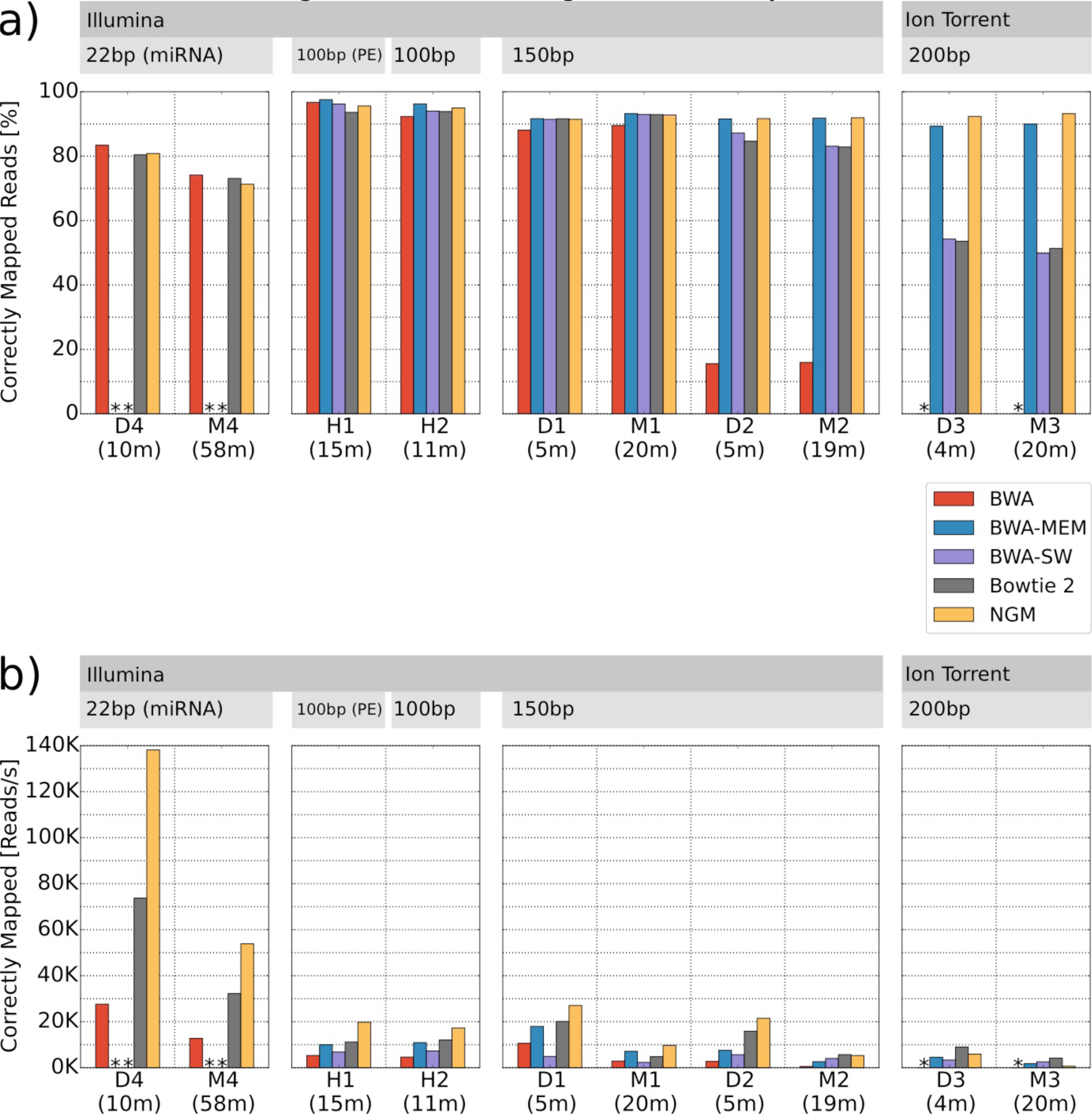
Mapping accuracy and mapping efficiency for different mappers and different input data. (a) Percentages of correctly mapped reads and (b) number of correctly mapped reads per second for five mappers. Table 1 gives a detailed description of the data. A * indicates that at least 99.9% of the reads were not mapped, due to limitations of the mapper. Teaser runtime in minutes is shown in parenthesis.

Next, we evaluated if BWA-MEM still performs best when using data from other model organisms or using different sequencing platforms and protocols. This evaluation is only possible with Teaser. Teaser required between 4 (D3) and 58 (M4) minutes to benchmark the data, running five mappers with their default parameters. The long runtime of M4 was mainly due to the runtime of BWA-MEM (41 minutes), which accounted for 70% of the total runtime. However we note that BWA-MEM was not designed for reads shorter than 70bp [12]. For the remaining data BWA-MEM used on average 17.9% of the total runtime.

For the *Drosophila melanogaster* (D1, D2) and *Mus musculus* (M1, M2) Illumina data sets, NextGenMap and BWA-MEM perform almost equally well in terms of correctly mapped reads (max. difference of 0.4%). For the simulated Ion Torrent data sets (D3, M3), NextGenMap showed a noticeably higher percentage of correctly mapped reads (3.02% and 3.23%, respectively) than BWA-MEM. For the miRNA data sets (D4, M4) BWA, and not BWA-MEM, had the highest rate of correctly mapped reads (83.41%, 74.15%). These results show the benefit of using Teaser to find the best mapper for a specific data set, especially considering that a 1% change in performance translates to tens of millions of additional reads mapped correctly in a genome-wide evaluation.

As highlighted by Fonseca et al. [3] and several others, runtime is crucial when selecting the best mapper especially in large-scale projects. In addition, for data sets like D1/D2 and M1/M2, where NextGenMap and BWA-MEM show very similar accuracies, runtime can be used to break the tie. To account for this, Teaser further reports the number of correctly mapped reads per second. Here, NextGenMap and Bowtie2 show the best performance, while BWA-MEM ranges from 2^nd^ to 5^th^ place depending on the data (Figure 2b).

### Automated parameter optimization and evaluation of mapping accuracy

Our previous results (Figure 2a) showed that BWA-MEM and NextGenMap performed best in terms of mapping reads to their correct position, while in terms of mapped reads per second (Figure 2b), NextGenMap and Bowtie2 outperformed the other mappers. However, the efficiency of Teaser with respect to computing times allows another application; Teaser can be used to identify parameter combinations that increase the fraction of correctly mapped reads and the number of correctly mapped reads per second.

**Figure 3:**
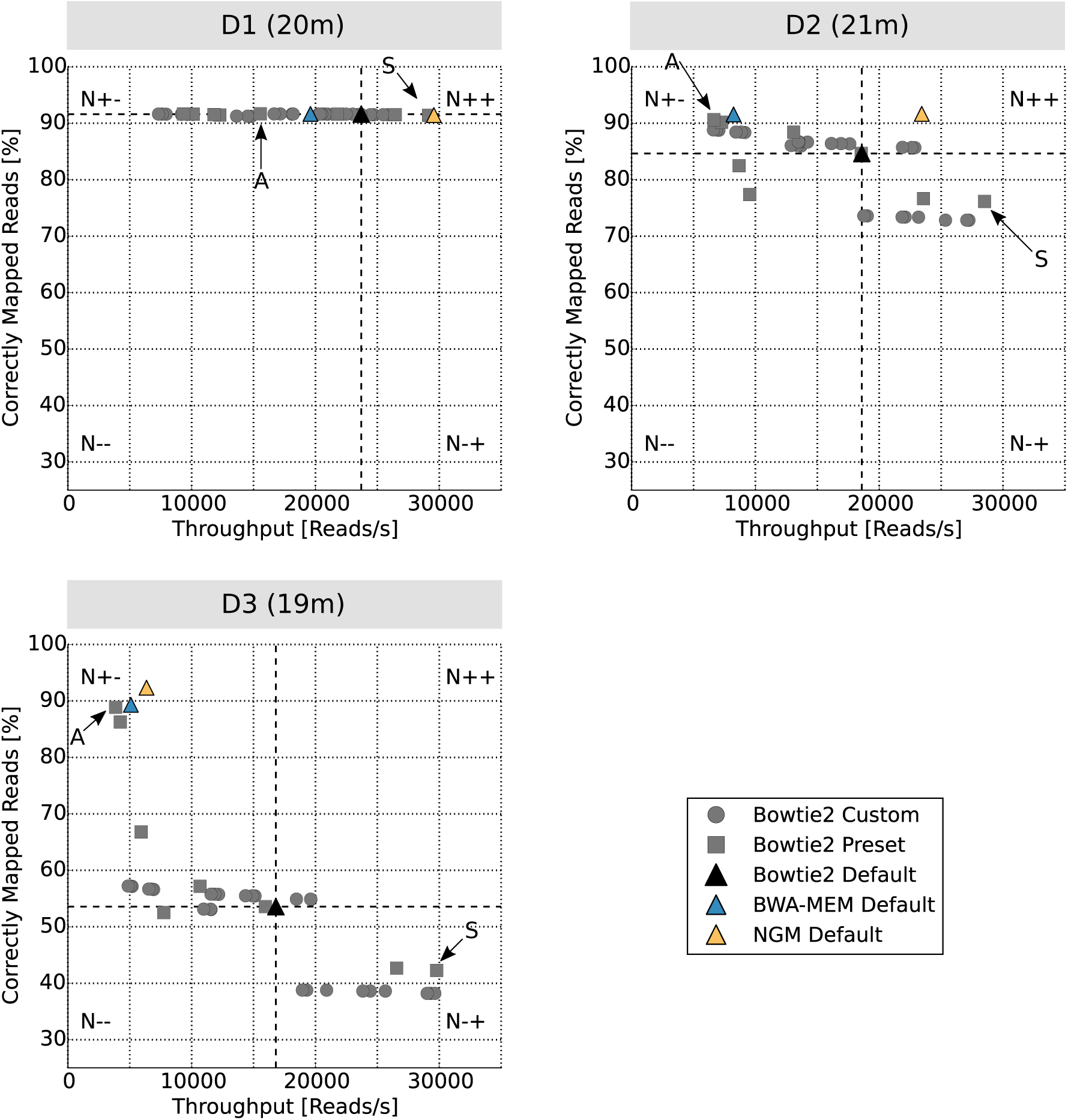
Throughput and percentage of correctly mapped reads for Bowtie2 when the mapping parameters were varied (grey circles and boxes). The black triangle shows the performance for the Bowtie2 default settings. While BWA-MEM and NextGenMap results using their default settings are shown as a blue triangle and a yellow triangle, respectively. Symbols above the dotted horizontal line show an increase in mapping accuracy if compared to Bowtie2 default, whereas symbols right of the dotted vertical line represent parameter settings with increased throughput. The number of parameter combinations that fell in each quadrant is specified by N_++_, N_+-_, N_--_, N_-+_ the “A” arrow points to the parameter combination that achieved the highest accuracy, whereas the “S” arrow marks the parameter combination with the highest throughput. The numbers in parenthesis behind the input data (D1, D2, D3) are the total runtimes needed to evaluate all parameter combinations on the respective data.

To show the versatility of Teaser, we ran Teaser for Bowtie2 with the default parameters, and eight parameter-options provided by Bowtie2 (e.g. “--very-fast” through “--very-sensitive”), the so-called Bowtie Preset parameters. In addition, we defined a custom range of values for three critical parameters: the length of the seed (-L), extending the alignment (-D), and the maximum number of times a repetitive read will be reseeded (-R). Thus a total of 34 different combinations of Bowtie2 parameters were evaluated and compared to BWA-MEM and NextGenMap, when the default parameters were used.

Figure 3 displays the number of correctly mapped reads per second (x-axis) and the percentage of correctly mapped reads (y-axis) for the 34 parameter combinations of Bowtie2 and the default parameters of BWA-MEM and NextGenMap for the *Drosophila melanogaster* genomic sequencing data (D1, D2, D3). For each of the three data sets the entire analysis finished in 21 minutes or less.

We see that changing the mapping parameters can lead to a large increase in the number of reads mapped per second and also in the fraction of correctly mapped reads. The plot can be divided in four quadrants relative to the position of Bowtie2 using the default parameter (black triangle): upper/lower by left/right. The lower left quadrant (N--) represents the parameter values that resulted in a lower amount of correctly mapped reads and a reduced throughput. Thus, they are outperformed by the default parameters. The lower right quadrant (N-+) encompasses the parameter settings that improve throughput, but correctly map fewer reads. The upper left quadrant (N+-) reflects parameter values that achieved a higher percentage of correctly mapped reads, but with reduced throughput. Those parameter combinations are interesting, but come at the expense of additional runtime. Thus, they may not be preferred in all applications. Finally, the upper right quadrant (N++) represents those parameter settings that outperformed the default parameter both in terms of speed and correctly mapped reads. Thus, the N++ parameter settings are always preferable to the default settings.

For the Drosophila resequencing experiment (D1) the default parameter values were substantially improved in terms of runtime (32% faster) at a marginal loss of 0.3% in accuracy. For the related species sequencing experiment (D2) and the Ion Torrent resequencing experiment (D3) some parameters settings were superior in terms of both speed and accuracy. For D2 we identified parameter settings that lead to a runtime improvement of 21% and that mapped an additional 1% of the reads correctly. The higher error rate and the length of the reads provided by Ion Torrent (D3) posed another challenge to optimize Bowtie2. Among the parameter combinations testes, Teaser found parameter values that led to a speed-up of 16% and a mapping accuracy that increased by 1.2%, compared to the Bowtie2 default values. More remarkably, one parameter combination (A-arrow Fig 3 D3) tested by teaser almost doubled the percentage of correctly mapped reads (from 53.3% to 88.6%), while achieving the same throughput as BWA-MEM (blue triangle).

Summarizing, systematically varying the parameter combinations of a particular mapper often leads to a substantial increase of accuracy and throughput. This optimization can be done within a few minutes and can be adapted to the specific data at hand. However, this analysis can be automatically extended to other mappers. Here we have only shown how to improve Bowtie2’s performance, but we could also easily attempt to optimize BWA-MEM, NextGenMap, or other mappers as well.

**Figure 4:**
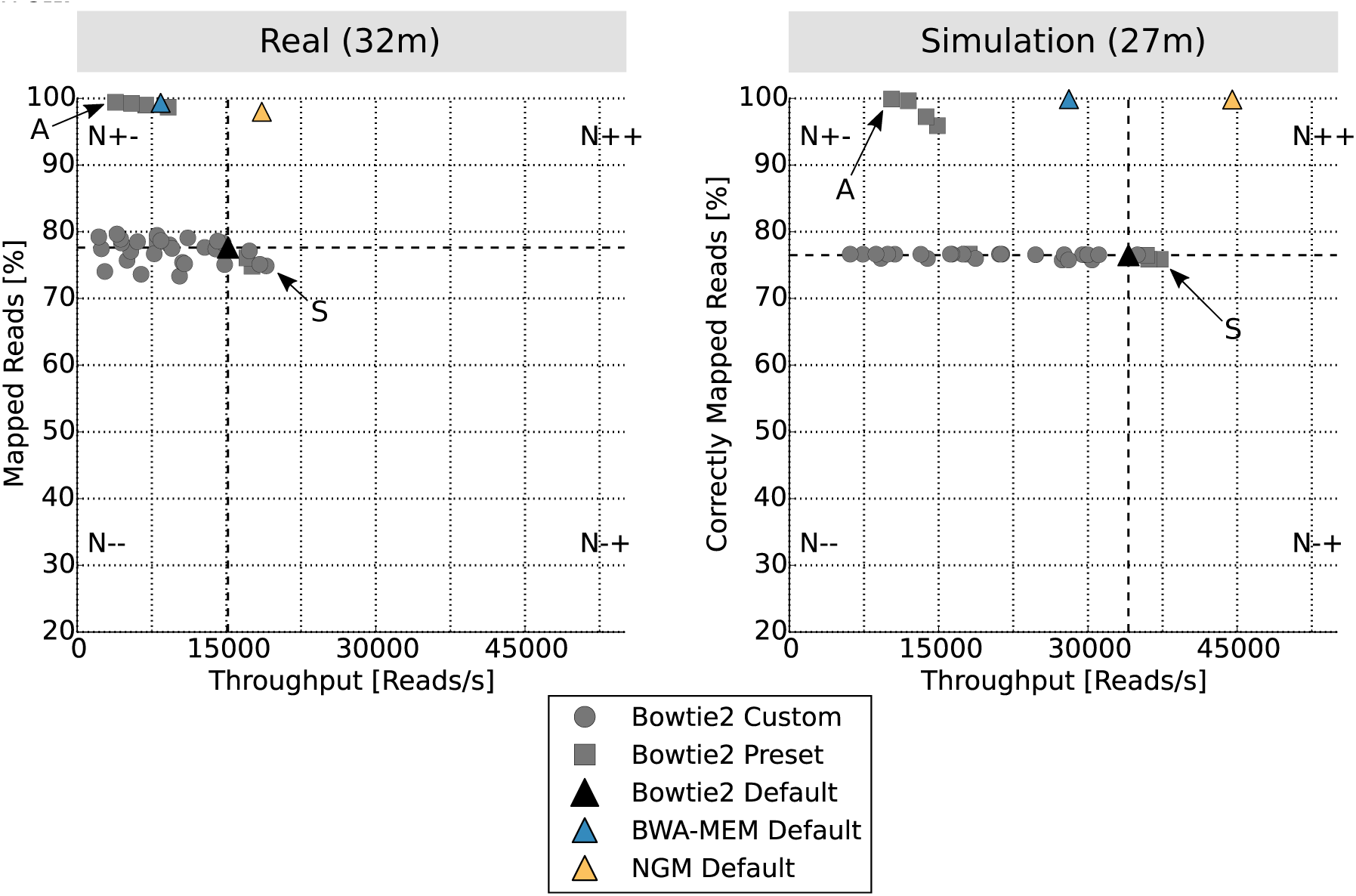
Benchmarking parameters on a real (a) and simulated (b) draft *de-novo* assembly. For the simulations Teaser sampled 100k reads of the *C. perifretum* data set and mapped it to the *de-novo* assembly of *C. rhenanus*. Simulation parameters were approximated based on a visual inspection of the real data set. For a detailed explanation of the legend we refer to Figure 3.

### Application to real data

Finally, we investigated how the mappers perform on a draft *de-novo* assembly using genuine and simulated Illumina data. We investigated the mapping of *Cottus perifretum* reads to the *de-novo assembly* of *Cottus rhenanus* (FJS, J.Cheng, J. <u>Altmüller, AvH, AW. Nolte</u>, under consideration). Teaser was used to map a subset of the real reads to the *de-novo* assembly using the same preset and default parameter settings of Bowtie2 and the default parameter settings of BWA-MEM and NextGenMap. Figure 4 shows the result for the parameter optimizations for real and simulated data. For real data, the true mapping positions are not available, so Teaser uses the percentage of mapped reads as the criteria to assess the performance of each run as described above. Teaser supports this statistic to allow a comparison of the optimized parameter settings and mappers for the real and simulated data. Thus recognizing if the parameters for the simulation were chosen reasonable or not. For example, if the simulated and genuine results strongly disagree, this can indicate if the genuine reads have unrecognized trimming and/or sequencing error issues.

Teaser automatically simulated and subsampled 100k reads for the simulated and real data, respectively. In both data sets the parameter settings of Bowtie2 perform equally well. Only the fastest parameter (indicated by S) changed for Bowtie2 from a preset set to a custom parameter defined by Teaser for the real data set. The overall throughput of the mappers and parameter settings changed between the simulation and the real data. This is expected to some degree as the simulation cannot fully mimic the complexity of the real data, especially when mapping to a *de-novo* assembly that may be incomplete and/or fragmented [16].

## Software

Teaser is available as a web application (teaser.cibiv.univie.ac.at), a virtual machine image and a command line version (https://github.com/Cibiv/Teaser) to increase the usability for expert and non-expert users. To further boost the applicability and advantage of using Teaser we provide different parameter settings used in this study on our github page (https://github.com/Cibiv/Teaser). Furthermore, we encourage the community to contribute parameter settings that improve the performance of mappers. We will incorporate such settings in the parameter files provided on the github page. Teaser is easy to use and at the same time extendable for other methods and parameters to be evaluated.

## Discussion

Choosing the most suitable read mapper and its parameters for a particular data set is far from trivial [3]. Improper mapper or parameter selection can result in many significant technical artifacts, including reduced coverage, reduced rates of heterozygosity, false determination of allele-specific expression, or other false results. Nonetheless, most current studies rely on default parameters of arbitrarily chosen mappers. Teaser seeks to overcome this deficiency by assisting in choosing the appropriate mapper and parameter setting by measuring the performance over an ensemble of different mappers and parameter combinations. This evaluation takes only minutes and does not require any manual intervention. Thus, Teaser is the first automated tool that finds and justifies the usage of a mapper and its parameter settings.

Our results for human data sets (H1, H2) are consistent with the results from GCAT [8]. At the same time, we demonstrate the necessity of benchmarking mappers individually for different combinations of sequencing methods, protocols, and genome complexities. For example, we find that BWA and not BWA-MEM to be a superior mapper for very short reads, and used Teaser to substantially improve Bowtie2’s mapping performance for three *Drosophila melanogaster* data sets and a real *de-novo* assembled data set. This effectively demonstrates the versatility and importance of Teaser, although such optimizations can and should be carried out for every mapper and for each experimental design. The analyses presented here only scratch the surface of Teaser’s potential, leaving the scientific community to fully explore Teaser’s full power.

From Figure 3 it is apparent that a simultaneous improvement of speed and accuracy is not frequently accomplished. However, it is often possible to improve accuracy with the expense of computing time or vice versa. Ultimately, the decision lies with the users whether they prefer parameter settings that are fast and accurate or more accurate but slower than the default. Teaser automatically provides such insights within minutes so that users can make an informed decision.

Teaser is easy to use and at the same time extendable to other methods and parameters combinations. Future work will include the incorporation of benchmarking RNA-Seq mappers and variant calling methods. We furthermore encourage the scientific community to contribute the optimal parameter combinations they detected to our github repository for their particular organism of interest. This will help others to quickly select the optimal combination of mapper and parameter values using Teaser.

## Abbreviations

NGM NextGenMap; NGS Next-Generation Sequencing

## Authors’ contributions

FJS, MS and PR conceived the method and designed the experiments. MS implemented Teaser. FJS, MCS and AvH wrote the manuscript. All authors read and approved the final manuscript.

## Acknowledgements

We would like to thank Han Fang for his helpful discussions. FS and MCS are supported through National Science Foundation awards (DBI-1350041 and IOS-1237880) and National Institutes of Health award (R01-HG006677). PR acknowledges support by the RNA-DK Biology (W1207-B09). AvH and MS acknowledge financial support from the University of Vienna and the Medical University Vienna.

